# Grasping the constraints of pure bacterial strains for the complete catabolism of micropollutants: a proteomic and kinetic study

**DOI:** 10.1101/2024.09.25.614793

**Authors:** Ana P. Lopez Gordillo, Alba Trueba-Santiso, Kilian E.C. Smith, Andreas Schäffer, Juan M. Lema

## Abstract

Research into the microbial degradation of organic micropollutants (OMP) often involves monitoring depletion of the parent compound and analyzing the biotransformation pathways that can lead to the production of metabolites, some being toxic, and/or to their mineralization. For the antibiotic sulfamethoxazole (SMX), previous studies testing a range of SMX concentration (mg down to ng L^−1^), have shown incomplete biotransformation of the parent SMX. This occurred both during wastewater treatment with mixed microbial communities and in studies with pure bacterial strains acclimated to SMX. This study explores the mechanism of SMX biotransformation and relationships with the proteome profile as possible reasons for the incomplete degradation of the parent SMX. *Microbacterium sp* BR1 served as an acclimated bacterial degrader of SMX in the range of µg L^−1^ to ng L^−1^. Depletion of the SMX was incomplete whereas the metabolite 3-amino-5-methylisoxazole (3A5MI) accumulated. The activity of the enzymes for the initial transformation of the parent SMX (SadA) was higher than that of further biotransformation steps (SadB). These results showcase that even a highly sensitive and metabolically active strain at very low SMX concentrations may require complementary enzymatic machineries to degrade metabolites that have an inhibitory impact in the biodegradation and persistence of this antibiotic.

**Synopsis:** A complete removal of organic micropollutants from water is challenging. This article delves into the bacterial degradation of the antibiotic SMX and proteome analysis to clarify underlying causes of its incomplete elimination

## 1. Introduction

Organic micropollutants (OMP) are a group of contaminants of major concern that are widespread in different environmental compartments. Despite their typically low concentrations in the environment of micrograms to nanograms per liter, they can represent a hazard for the environment and human health. Biological treatment, such as microbial biotransformation, are widely used for OMP conversion into less dangerous compounds, and if possible their mineralization. The latter is their complete degradation^1^ accompanied by the release of CO_2_.^2^ In some cases, however, biotransformation results in the formation of metabolites which can be even more toxic and less biodegradable than the original parent OMP molecule. Consequently, investigations into the biotransformation of OMPs should include quantification of any depletion in the parent OMP supplemented by analysis of the formed metabolites.^3^

Numerous studies into the general environmental fate of legacy compounds, such as chlorinated solvents or hydrocarbons^2,4^ exist. However, studies into the actual biotransformation mechanisms behind OMP removal are more recent and these include pharmaceuticals, industrial chemicals and pesticides.^1^ Among these, antibiotics are of particular interest due to their potential to unleash the development of bacterial resistance mechanisms.^5,6^ From within this group, the sulfonamides have been widely investigated^7^, with sulfamethazine (SMZ), sulfadiazine (SDZ), sulfapyridine (SPY) and sulfamethozaxole (SMX) being among the most commonly investigated compounds.^8^

Four main processes for sulfonamide breakdown have been identified for mixed bacterial strains or axenic cultures^8^: 1) cleavage of the heterocyclic ring at the N-O bond, 2) cleavage of the sulfonamide bond (S-N), 3) cleavage of the C-N bond, and 4) cleavage of the aniline moiety. These processes may occur separately or combined which leads to a wide diversity of possible metabolites. Furthermore, the occurrence of co-metabolism or direct metabolism also leads to formation of diverse transformation products.

SMX direct metabolism involves breakdown of the parent compound, while its co-metabolism may involve simple reactions that only modify the parent compound and in fact enhance or partially keep its toxicity.^9^ Some single transformation reactions accomplished co-metabolically by bacteria in soil and water, as well as in axenic cultures^8^ in the presence of SMX include acetylation, hydroxylation or nitration at the *para* amino group. Various reported biotransformation products produced co-metabolically include N^4^-acetylsulfamethoxazole (Ac-SMX)^10^, N^4^-hydroxy-acteylsulfamethoxazole (OH-Ac-SMX)^11^, 4-hydroxyl-N-(5-methyl-1,2-oxazole-3-yl)benzene-1-sulfonamide (4-OH-SMX)^12^ and 4-nitro-sulfamethoxazole (NO_2_-SMX).^13^ Direct metabolism of SMX originates a wider range of metabolites of which some examples include: 4-aminophenol, hydroquinone, 3-amino-5-methylisoxazole (3A5MI), 1,2,4-trihydroxybenzene, 4-aminothiophenol, 4-amino-benzesulfonic, 4-amino-benzesulfonamide, aniline and sulfanilamide.^14–18^

Sometimes SMX and its formed metabolites may have an incomplete biological removal that impedes achieving mineralization. Possible causes for this may be linked to the toxicity of the formed metabolites and their back transformation to the parent OMP (retro-inhibition).^19,20^ Some examples include the metabolites NO_2_-SMX and 4-OH-SMX which are toxic and produced an inhibition effect on *Vibrio fischeri* growth.^9^ Also some SMX metabolites from axenic cultures, which are initially stable, can be back transformed to SMX when present in complex environments.^21^ Such back transformation has been described for Ac-SMX^22^ and 4-NO_2_-SMX, the latter under nitrate starvation.^23^

Proteomics is a tool that allows a deeper understanding of the (in)complete degradation of OMP, as it provides information on the degradation mechanism of xenobiotic removal and on the bacterial metabolic response^24^ as an effect of the exposure to a pollutant. Consequently, the proteome analysis may include functional proteins usually involved in the general bacterial metabolism and/or proteins specific to the catabolism of OMP of interest. Studies on SMX biodegradation may include the analysis of proteins and resistance mechanisms involved in sulfonamides biodegradation, such as: sulfonamide monooxygenases (Sad cluster)^25^, modifications of the enzyme dihydropteroate synthase (DHPS) (*fol*P and sul genes: *sul* 1-*sul*4)^26^, enzyme arylamine N-acetyltransferase (NAT)^27,28^ and multidrogue efflux pumps systems (MexAB-OprM and *smeDEF)*.^29,30^

In general, the biotransformation of OMP is influenced by their bioavailable concentration, the microbial community, its metabolic activity and the environmental conditions^2^. Although several investigations into SMX biotransformation have been done at mg L^−1^ concentrations, the underlying mechanisms may differ at lower concentrations which are more representative of those found in the environment. Therefore, research at the low µg L^−1^ level close to the ng L^−1^ become of relevance.

An study focusing on SMX bacterial biodegradation at low concentrations emphasized on existing constraints for achieving mineralization. Although it was shown that adapted degraders are able to metabolize low SMX concentrations, an incomplete removal was described.^31^ In the present study we used *Microbacterium* sp. strain BR1 as a degrader model to investigate the biotransformation mechanisms of SMX at different low concentrations, and to elucidate the reasons why a complete SMX removal is not achieved. For this goal we combined analytical chemistry (liquid chromatography and mass spectrometry) and mass spectrometry-based proteomics.

## 2. Materials and methods

### 2.1 SMX solutions

SMX (purity 98%,Sigma Aldrich) was used to prepare a stock solution of 20 mg L^−1^ in Milli Q water. Through dilutions three solutions of SMX were prepared in phosphate saline buffer (PBS) pH 7.4 and sterilized at 121°C for 15 minutes. These solutions were spiked to the corresponding reactors to test the biotransformation of three concentrations of SMX: 20 µg L^−1^, 12 µg L^−1^ and 0.1 µg L^−1^.

### 2.2 Biomass acclimation and production

*Microbacterium sp* BR1 was selected as a suitable bacterial degrader of SMX. A sample of this pure strain was provided by Dr. Boris Kolvenbach from the Institute for Ecopreneurship, University of Applied Sciences and Arts (Northwestern Switzerland). Sterile standard media 1 at 25% (Carl Roth) containing 1mM SMX was used for the cultures.^32^ Culture flasks were kept in darkness and incubated at 28 °C with an agitation of 140 rpm. When the optical density (OD_600_) reached a value of 1.4, the cultures were centrifuged at 7000 g and 4 °C for 20 min and washed with cold sodium chloride (NaCl) 0.85 %. After two washes, the bacterial pellets were homogeneously resuspended in NaCl 0.85 % and glycerol 20 %. The resuspension was split in several aliquots and stored at −80°C until their use in the biotransformation tests. The acclimation to SMX aimed to trigger the enzymes involved in the catabolism, which is necessary in such short biotransformation tests.

### 2.3. Biotransformation tests

One batch experiment was performed per SMX test concentration. In each batch, the reactors consisted of 50 mL amber glass bottles filled with 26 mL of sterile PBS. To obtain the required initial SMX concentration, 1 mL of the sterile SMX spiking solution was added to each bottle. Biotransformation assays were started by the addition of 1 mL of thawed *Microbacterium* sp BR1 aliquots to each reactor. The bacterial density used to start the experiments was determined by streaking serial dilutions of the aliquot on agar plates and calculating the colony forming units (CFU). This was 209.8 ± 0.34 x 10^6^ CFU mL^−1^. To exactly reproduce the conditions in the parallel tests for measuring SMX mineralization^31^, an insert preloaded with 1 M potassium hydroxide (KOH) was added to the bottles before their closure with a screw cap. Each reactor comprised biological triplicates. Negative controls consisted of bottles without bacteria. The reactors were kept at 22 °C under horizontal agitation at 140 rpm in the darkness.

Each biotransformation test lasted 24 hours. During this period, four sampling points were defined: 2 h, 4 h, 8 h and 24 h. For each time point, triplicate reactors from each test concentration were sacrificed and opened inside the sterile bench, the KOH and the insert were carefully withdrawn and disposed (these were only added to reproduce the set-up used for the mineralization experiment). The reaction medium was mixed before transferring it to 50 mL sterile tubes and centrifuging at 2190 g and 4°C for 35 minutes. The supernatant was separated from the bacterial pellet, frozen and stored for further analysis of SMX and the metabolite 3A5MI using Liquid Chromatography coupled to Mass Spectrometry (LC-MS/MS). The bacterial pellet was washed twice with cold NaCl 0.85%, centrifuging using the same conditions as above and the supernatant discarded each time. The washing procedure was repeated once. Afterwards, the bacterial cells were resuspended in 2 mL of cold NaCl 0.85% and frozen until their preparation for proteome analysis.

### 2.4. Analytics of SMX and 3A5MI

Defrosted supernatants of the test at 20 µg L^−1^ and 12 µg L^−1^ were centrifuged again at 5974 g and 4° for 20 minutes. An SMX standard addition approach was used for the centrifuged samples to counteract any matrix effects. A 100 µL volume of sample was injected into an ultra-high-performance liquid chromatograph (UHPLC ELUTE, Bruker). Sample up-concentration was achieved via online extraction (OLE) prior to the chromatographic separation in a C18 column (Intensity solo, Bruker). In the mass spectrometer (timsTOF PRO, Bruker), the analytes were ionised with electrospray ionization (ESI) in positive mode and fragmented with a broad band collision-induced dissociation (bbCID). For the screening, a triple quadrupole (QQQ) and a Quadrupole Time-of-Flight (QTOF) analyzers were used. Details on the analytical method are given in the Supplementary information (Text S1 and Table S1). Spiking solutions and filtered inocula were analyzed for chemical contamination. Measurement of the supernatants from the 0.1 µg L^−1^ test were below the analytical limit of detection. Moreover, the detection and quantification of additional metabolites were not feasible to difficulties with the analytic methodologies.

### 2.5 Proteome analysis

Samples from the 20 µg L^−1^, 12 µg L^−1^ and 0.1 µg L^−1^ reactors were pretreated as follows: one bacterial pellet replicate from each sampling point was thawed and centrifuged at 5974 g and 4°C for 20 minutes, followed by the disposal of the supernatant (0.85% NaCl). Proteome extraction was done according to the method described in Kennes et al study.^33^ For this, protein release and denaturalization was achieved through cell lysis using digestion with 1% of sodium dodecyl sulfate (SDS) pH 7.5 at 90°C for 20 minutes and four cycles of mechanical disruption with glass beads beating each with a duration of 3 minutes. After centrifugation of 1218 g at 4°C for 20 minutes, the proteins contained in the supernatant were transferred to microcentrifuge tubes for their precipitation with cold acetone and incubation at −20°C. Acetone was removed after centrifugation at 10621 g and 4°C, and the proteins resuspended in molecular grade water and acetone (1 volume per 4 volumes). An incubation at −20°C with fresh acetone was repeated and the supernatant discarded. Next, the extracted proteins were resuspended in water molecular grade and frozen.

Protein quantification of the extracts was determined using the Pierce bicinchoninic acid protein assay kit (BCA, Thermo Scientific) (Table S2). Additionally, an SDS-PAGE electrophoresis with NuPAGE gel (4-12% Bis-Tris acrylamide, Thermo Fischer) was performed under denaturing conditions to verify any protein degradation in the extracts. The electrophoresis was run with aliquots containing 10 µg protein and the gel stained with a standard Coomassie protocol (Figure S1).

For the mass spectrometry analysis, the protein extracts were trypsin digested and desalted. The peptides from the mixture were injected to a nanoELUTE chromatograph (Bruker) with an Aurora analytical column (C18, 250 × 0.075 mm, 1.6 μm, 120 Å, IonOpticks).^34^ The nHPLC was configured with binary mobile phases that included solvent A (0.1 % formic acid in miliQ H_2_O) and solvent B (0.1 % formic acid in acetonitrile). The analysis time was 105 min, in which the B/A solvent ratio was gradually increased. Blanks were injected between samples with an analysis time of 60 min, confirming no carry-over. For mass spectrometry (MS) acquisition, a collision-induced dissociation (CID) fragmentation and a nanoESI positive ionization mode was employed. PASEF-MSMS scan mode was established for an acquisition range of 100–1700 m/z.^35^ MS analyses were performed at the Mass Spectrometry and Proteomics Unit (Area of Infrastructures) of the University of Santiago de Compostela. Peptide identification was done with the software tool PEAKS Studio (Bioinformatics Solutions, Canada) and compared to the private genomic database of the strain *Microbacterium sp* BR1 provided by Dr. Boris Kolvenbach from the Institute for Ecopreneurship, University of Applied Sciences and Arts (Northwestern Switzerland).

Analysis of the peptides was done with a label-free semi quantification based on the spectral counting method and the Spec value.^34^ Spec values were considered as the relative abundance of the enzymes linked to SMX metabolism by *Microbacterium sp* BR1 (Sad cluster and DHPS). Additional bioinformatic analysis included processing the Gene Ontology (GO): Biological process (BP) and Molecular function (MF) with the Unipept Desktop 3.0 software^36^ to investigate the effect of the tested SMX concentrations on diverse functional proteins. GO containing less than 2 peptides were neglected for the analysis.^37^

## 3 Results and discussion

### 3.1. Biotransformation of SMX by *Microbacterium sp BR1*

It has been previously reported that *Microbacterium sp* BR1 is able to mineralize up to 60% of parent SMX when provided as the sole source of carbon and energy at environmentally relevant concentrations (25 µg L^−1^ down to 0.1 µg L^−1^).^31^ In the present study, the analytical measurement of the reaction media intended to delve into the biotransformation of SMX occurring in a concentration range comparable to the reported for mineralization.

The commencement of the catabolic pathway of SMX by strain BR1 and some of the expected initial metabolites are shown in Figure 1. The catalytic flavin monooxygenase (Sad A) together with the flavin reductase (Sad C) are responsible for the initial attack of sulfonamide molecule resulting in the release of 4-benzoquinone imine (BQI), 3A5MI and Sulfur dioxide. In the catabolic pathway reported for *Microbacterium sp* BR1, BQI derive into Benzoquinone and 4-aminophenol ^32,38^, whereas 3A5MI is predicted to generate the metabolite (3-aminoisoxazol-5-yl) methanol under aerobic conditions (pathway prediction system: EAWAG-BBD)^39^, such as the ones present in our study.

**Figure 1.**
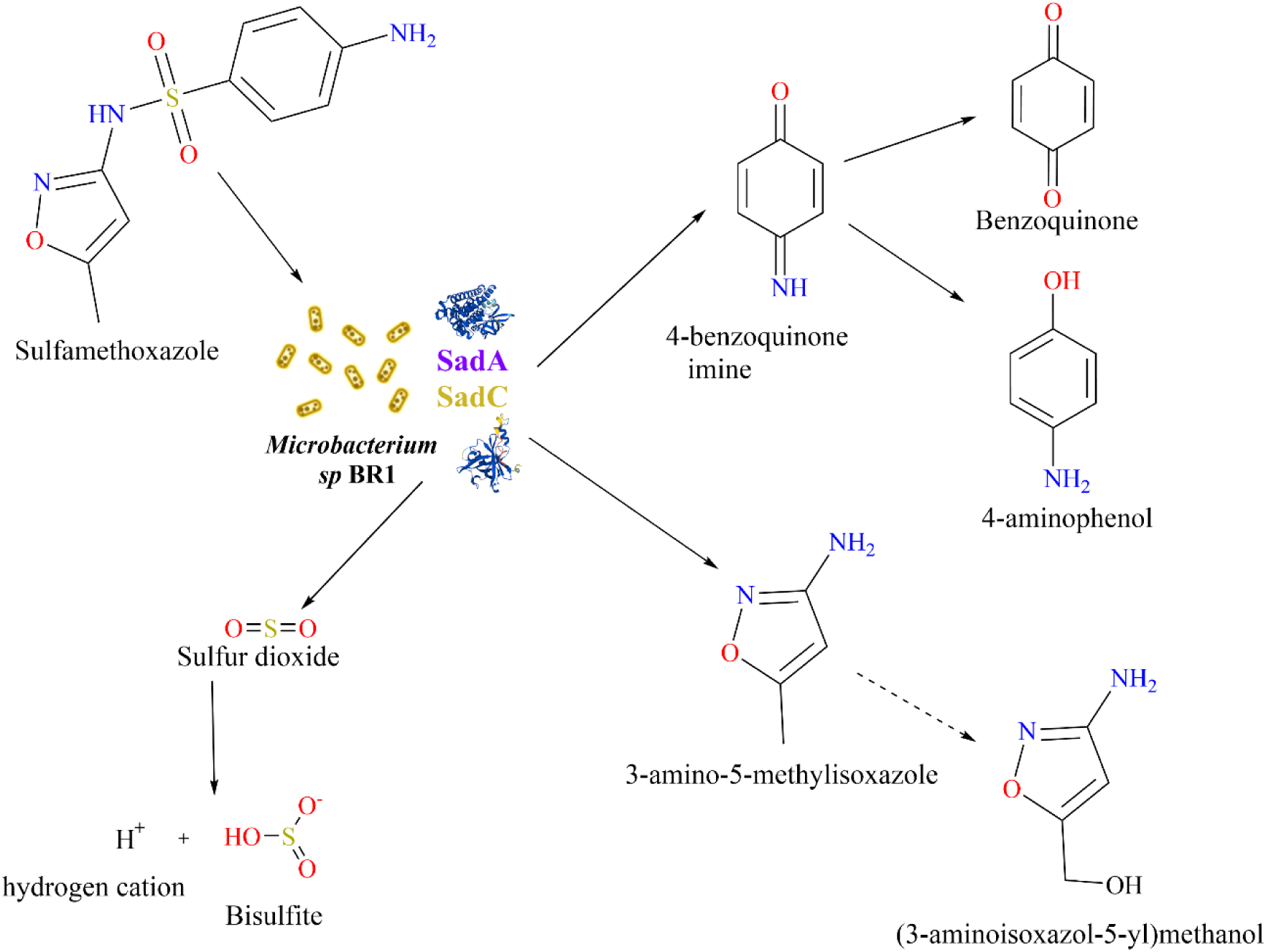
The ipso-hydroxylation has been described as the first step in SMX catabolism by *Microbacterium sp* BR1. After this step, the initial SMX molecule is split into 3A5MI and 4-benzoquinone imine. The solid arrows indicate the pathway reported in literature^32,38^ while the dotted arrow points to the predicted pathway.^39^

In our study, the analytical measurement of the supernatant endorsed that the depletion of parent SMX coupled the production and accumulation of the metabolite 3A5MI in the test of 20µg L^−1^ and 12 µg L^−1^ (Figure 2 and Figure S2 accordingly). The initial catabolic step of the ipso-hydroxylation of the parent SMX led to a pronounced drop at the 2 h of the test. After this, the residual SMX reached a plateau that kept on over time, akin to the reported during the mineralization of SMX in a similar concentration (25 µg L^−1^).^31^ This is also in close agreement with previous investigations performed with much higher concentration (25 mg L^−1^).^40^, which suggests that the catabolism performs comparably in a wide range of SMX concentrations.

**Figure 2.**
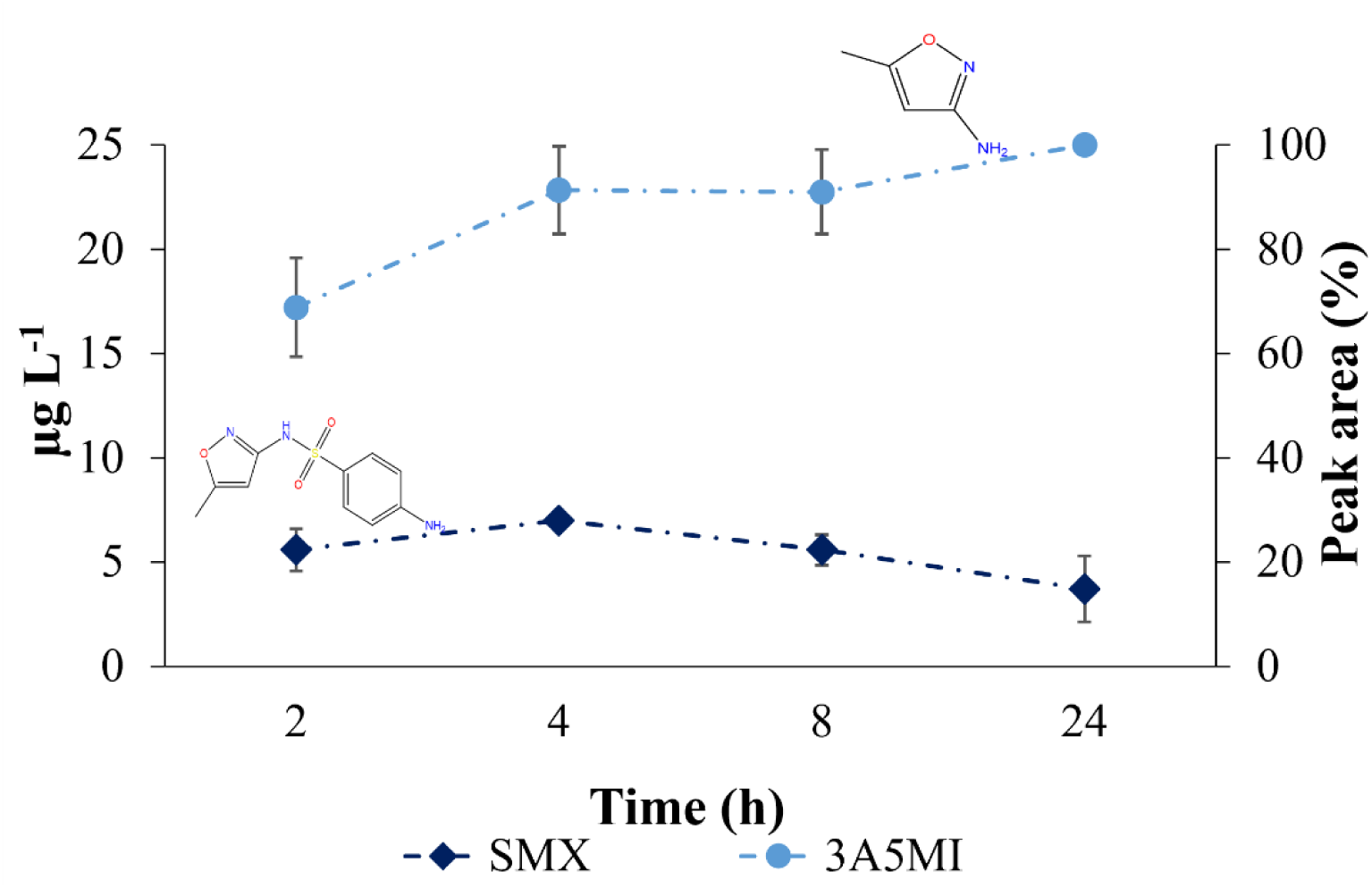
Evolution of SMX (diamond) and 3A5MI (circle) during biotransformation test with an initial concentration of 20 µg L^−1^. 3A5MI corresponds to a relative detected peak area. Depicted values are means of triplicates with their standard deviations.

The plateau of residual SMX over time regardless of the different initial SMX concentrations and in the presence of an acclimated bacterial fostered the investigation of the proteome profile to reveal a source of the incomplete biotransformation and the described incomplete mineralization (60% of parent SMX).^31^

### 3.2. Proteome expression of *Microbacterium* BR1 throughout the SMX biotransformation experiments

SMX biodegradation studies^25,38^ report that the catabolism of the sulfonamides rely on the key catalytic enzymes designated Sad ABC cluster. After the initial step facilitated by Sad A, the production of 4-benzoquinone imine primes the path towards mineralization and concomitant production of biomass (Figure 3). The formed 4-aminophenol is further transformed into 1,2,4-trihydroxybenzene by the flavinprotein monooxygenase SadB alongside SadC, which provides the necessary reduced FMN^38^. As for the path of 3A5MI, it has been described as dead end for *Microbacterium sp* BR1.

**Figure 3.**
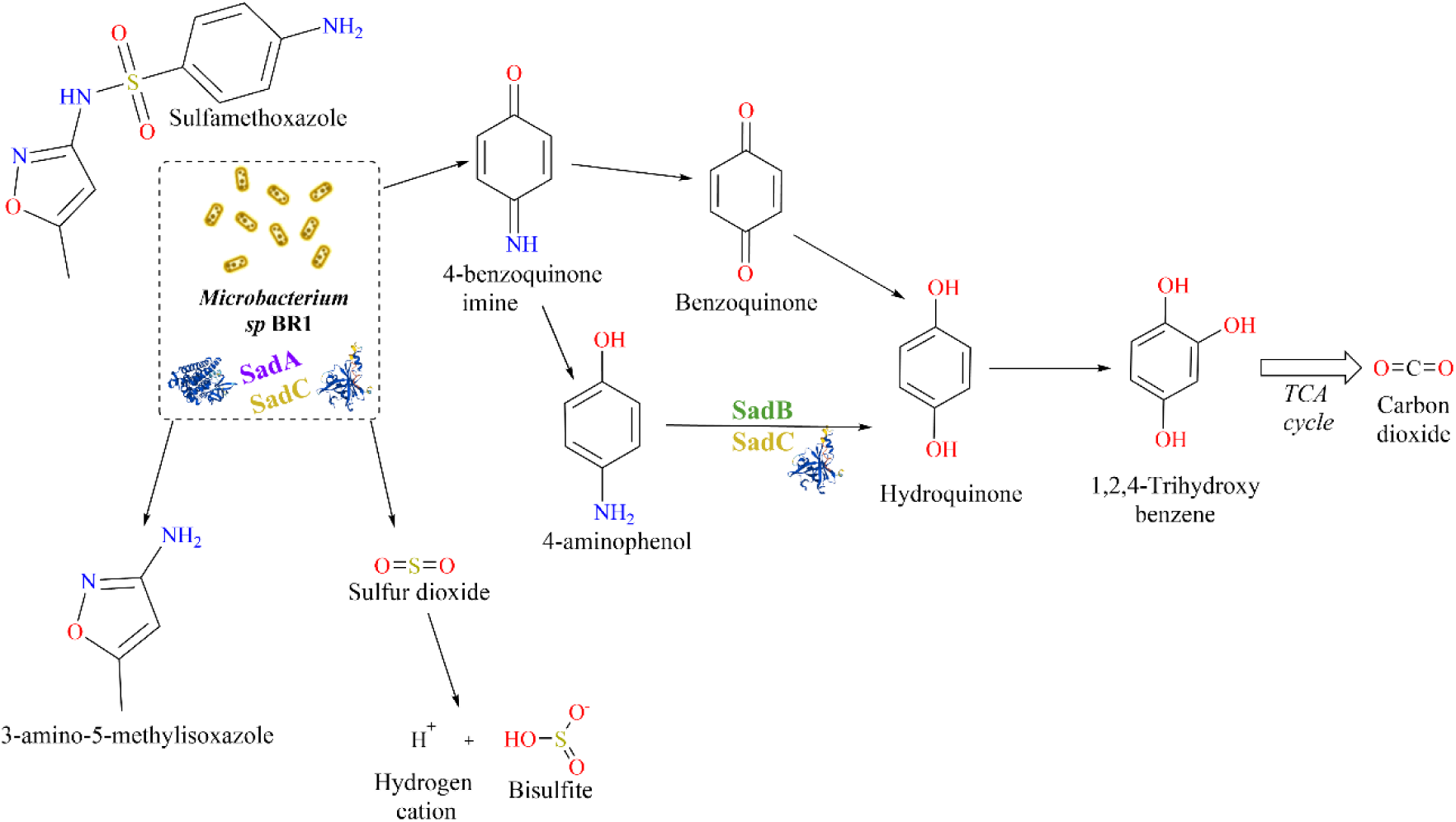
Reported metabolic pathway of the antibiotic ^14^C-SMX by *Microbacterium sp* BR1.^32,38,40^ The key enzyme cluster SadABC is displayed in the corresponding catabolic step. Mineralization could be tracked in the form of radioactive ^14^C-CO_2_ and linked to the degradation of the commonly radiolabeled ring phenyl.^31^ Metabolic pathway modified from original scheme.^38^

To better understand the impediments to achieve SMX complete metabolization, we determined the expression of the different subunits of the SadABC protein complex (sulfonamide degrading enzymes) over the course of the experiments (Tables S3-S8, Figure 4, A-C). The catabolic activity of *Microbacterium sp* BR1 was triggered in all the cases detecting an increase on the abundance of SadA and SadB over the time in those experiments with 12 and 20 µg SMX L^−1^. It was particularly interesting to detect this triggering effect even at an initial concentration of 0.1 µg L^−1^ of SMX. However, in this case, there was a further drop that could be associated to SMX nearly exhaustion. Differently to the other Sad complex units, SadC expression was constant during the time in all the concentrations investigated (Figure 4).

**Figure 4.**
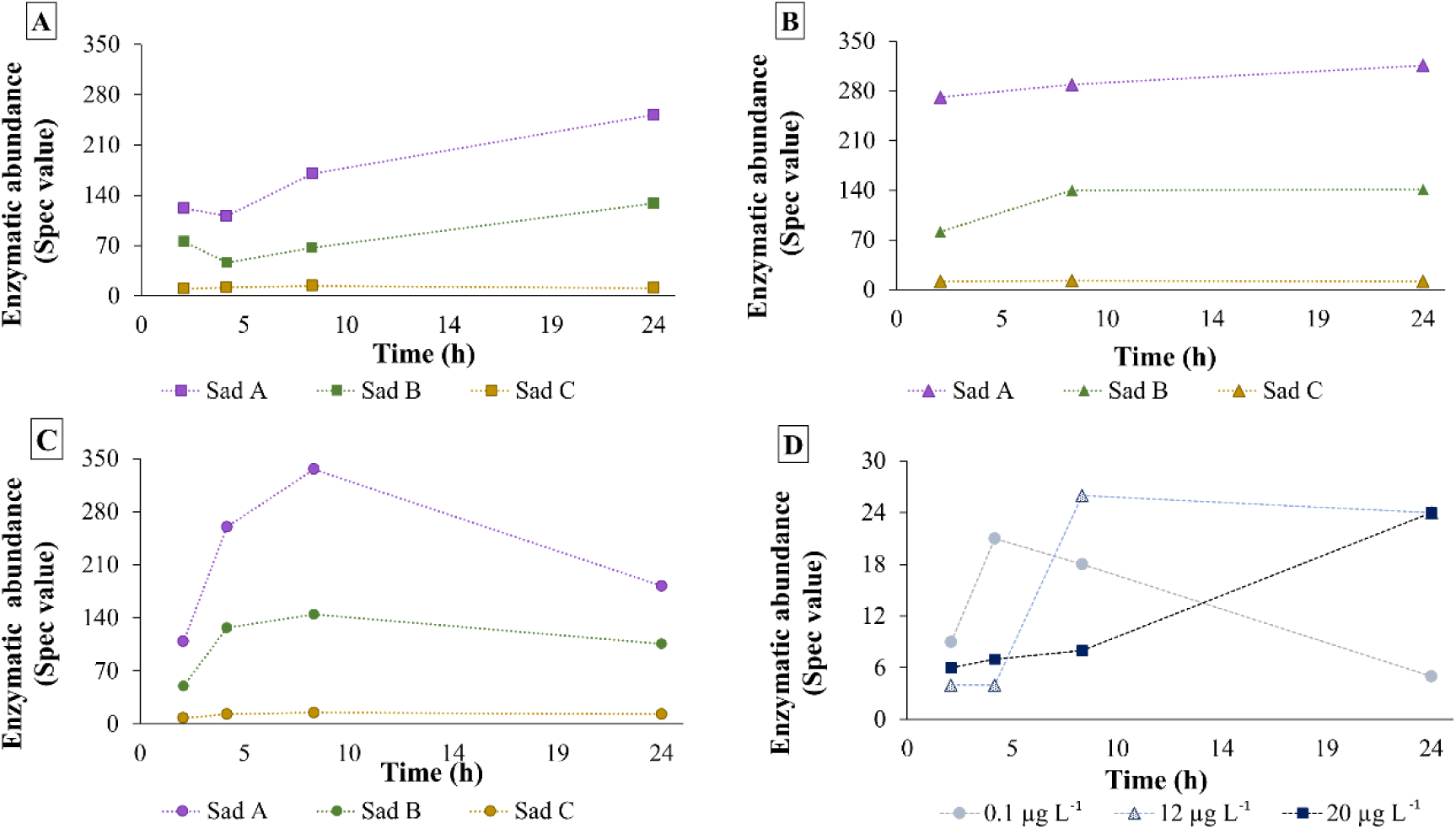
Relative abundance of Sad enzymes cluster and Sul1 over time in tests with different SMX initial concentrations: A) 20 µg L^−1^, B) 12 µg L^−1^ and C) 0.1 µg L^−1^. Board D depicts the relative abundance of Sul1 in the three test SMX concentrations.

SMX acts as a competitive inhibitor of the enzyme DHPS, which is involved in the production of the folate precursor necessary for the bacteria to reproduce.^41,42^ Some resistance mechanisms that allow bacteria to withstand this effect of sulfonamides include target modification (i.e.gene mutations) related to DHPS (*fol*P and *sul*1-*sul*4) and multidrug efflux systems.^26,29,30,43^ The sulfonamide-resistant DHPS gene, *sul*, encodes for a variation of this enzyme with low affinity to sulfonamides. Different SA-degrading strains (e.g. *Actinobacteria* or *Micrococcaceae*) with known genome sequences contained both *sad* and *sul* genetic clusters, which originated the hypothesis that sulfonamide degradation might be dependent on sulfonamide resistance.^44^ In our experiments, we detected the expression of Sul in all three concentrations (Figure 4, D). At the highest concentration (20 µg L^−1^), there was a clear trend of expression increase over the time, while in 12 µg L^−1^ there was a plateau and at 0.1 µg L^−1^ there was an initial increase followed by a decay. The fact that Sad and Sul followed similar expression trends throughout the time at the different concentrations tested could strengthen the hypothesis of sulfonamide resistance and degradation being associated in this strain and providing biochemical evidence for it.

The peptides detected in our samples from the 12 µg SMX L^−1^ experiments were categorized according to the Gene Ontology classification of molecular functions^45^ (Tables S9-S10). This allowed to detect increases when comparing the samples from 24 h to 2 h in the categories cell division and cell cycle, in general cellular maintenance activities such as phosphorylation, translation, or DNA replication (Figure 5). Moreover, the carbohydrate metabolic process and the tricarboxylic acid cycle also increased confirming that bacteria were assimilating carbon during all the duration of the experiments. These proteome expressions indicate that at the end of the experiment *Microbacterium* cells were still active, and therefore, the biotransformation of the parent compound and transformation products stopped due to other reasons.

**Figure 5.**
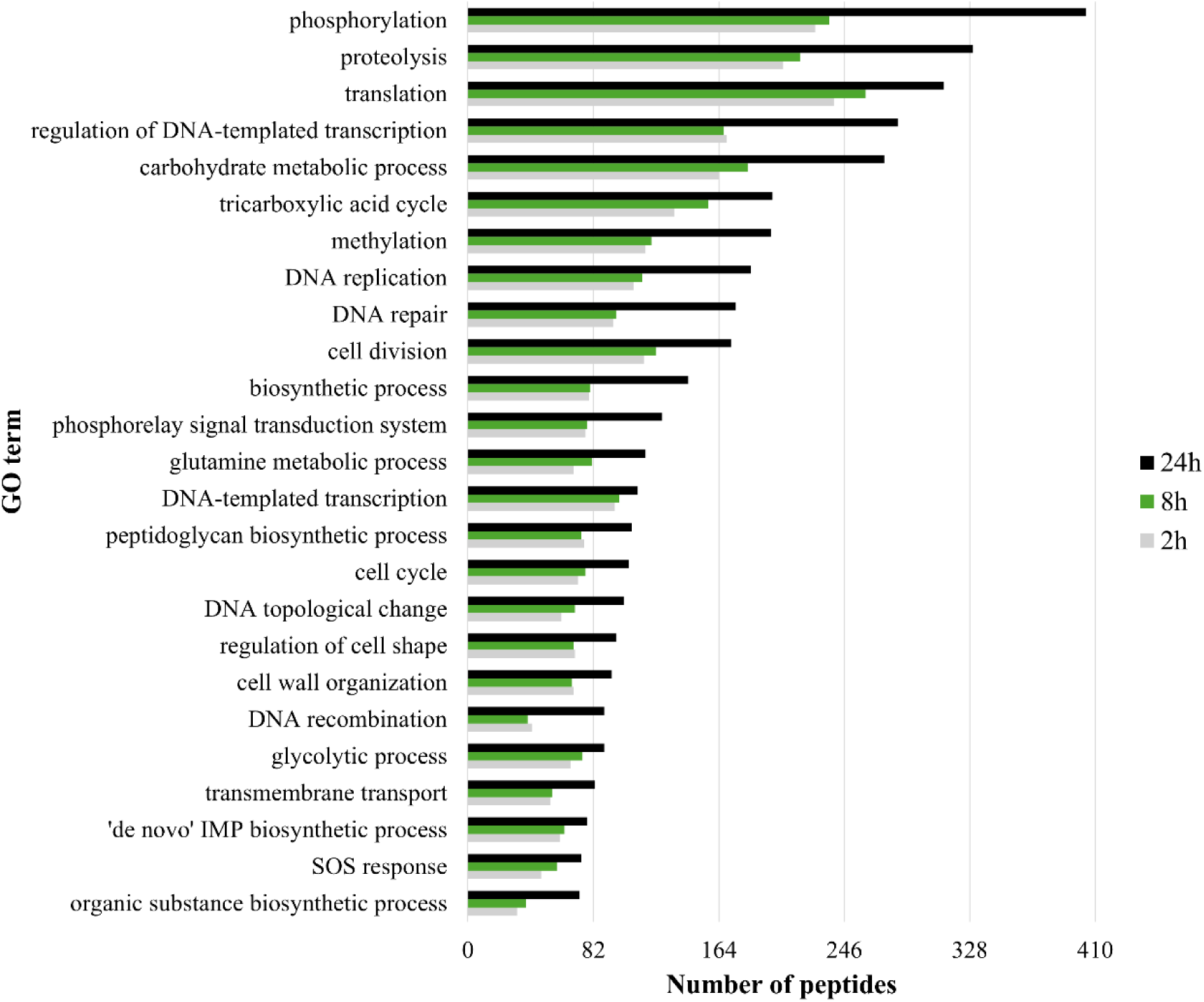
Number of peptides belonging to selected 25 Gene Ontology biological processes categories that increased its expression over the time in those experiments performed with 12 µg SMX L^−1^.

Interestingly, an increase in peptides related to regulation of cell shape and cell wall organization (Figure 5) occurred over the time which could be attributed to morphological changes on the cells as a mechanism of antibiotic survival. Previous reports on the literature pointed to changes in bacterial morphology as a consequence of exposure to antibiotics.^46^

Enzymes previously found in different microorganisms which enable aromatic ipso-substitutions and decreases the susceptibility towards antibiotics include laccases^47^, versatile peroxidases (VP)^48^, 4-sulphobenzoate 3,4-dioxygenase (PSB dioxygenase system)^49^, monooxygenases (CYP3A4, CYP3A5, CYP2D6*1)^50^ or the flavoprotein monooxygenases 6-hydroxy-3-succinoyl-pyridine (HSP) hydroxylase (HspB).^51^ However, these enzymes are not encoded in the genome of *Microbacterium sp* BR1^44^ and therefore, not present in our samples.

### 3.3. Kinetics of SMX: inhibition due to accumulation

As explained in the above sections, *Microbacterium sp* BR1 did not show limitations on its enzymatic activity nor its viability. Hence those can be excluded as potential explanations for the incomplete removal of SMX during the biotransformation tests. The paragraphs below address additional considerations related to the incomplete removal of SMX.

As the relative abundance of catalytic enzymes (Sad) and the functional proteins related to cell growth, carbon utilization and cell maintenance raised (Figure 4 and Figure 5), it can be presumed that the produced metabolites (including 3A5MI) are not toxic to the strain BR1. This aligns to the statement mentioning that 3A5MI lacks any toxicity.^9^ A similar scenario was reported for *Achromobacter denitrificans* strain PR1, another SMX bacterial degrader that produces and accumulates 3A5MI as main metabolite lacking a toxic effect.^52^

Some pure bacterial strains were reported capable of cleaving SMX oxazole ring and 3A5MI ^18,53,54^ yielding to different metabolites than the observed for *Microbacterium sp* BR1, which shows that the degradation mechanisms vary even between sulfonamide degraders. Impediments on SMX removal in pure strains can be overcome in mixed cultures, where the different enzymatic machineries become complementary.^55^ This has been reported for 3A5MI further catabolized in mixed cultures.^56^ Considering the previous, *Microbacterium sp* BR1 presumably does not possess the enzymatic machinery to degrade 3A5MI and it accumulates consequently.

3A5MI accumulated parallelly to the deceleration in the biotransformation of the parent SMX, which became negligible from 2 h onwards (Figure 2). A delayed biotransformation has also been referred previously, where the biodegradation rate of parent SMX decreased by 28% due to the presence of 3A5MI.^15^ Since 3A5MI has not been reported to have a back conversion to parent SMX, the retro-inhibition is excluded of triggering the partial removal of SMX. Instead, a substrate inhibition phenomenon seems to be a plausible explanation for the constant residual SMX observed in Figure 2.

The findings of this study contribute to deciphering the biotransformation mechanism causing the incomplete biodegradation of SMX and can serve as a basis to explain the systematic incomplete biotransformation of organic micropollutants. In this sense, future research could focus on enhancing OMP degradation with mixed cultures of selected degraders known to bear complementary degradation pathways.

## Acknowledgements

We especially thank Prof. Philippe Francois-Xavier Corvini and his team for kindly supplying the *Microbacterium* sp. BR1 which allowed us to perform this study. This work was funded by the European Union’s Horizon 2020 research and innovation program under the Marie Skłodowska-Curie grant agreement No. 812880 (MSCA-ITN-2018: EJD Nowelties). Alba Trueba-Santiso acknowledges a Juan de la Cierva-Formación postdoctoral grant (FJC2019-041664-I). Alba Trueba-Santiso and Juan M.L. belong to the Galician Competitive Research Group (GRC)_ ED431C-2021/37. Authors would like to thank the use of Mass Spectrometry and Proteomics facilities (RIAIDT-USC analytical facilities) from Santiago de Compostela University.

## Notes

The authors declare no competing financial interest.

---

**Figure S1.**
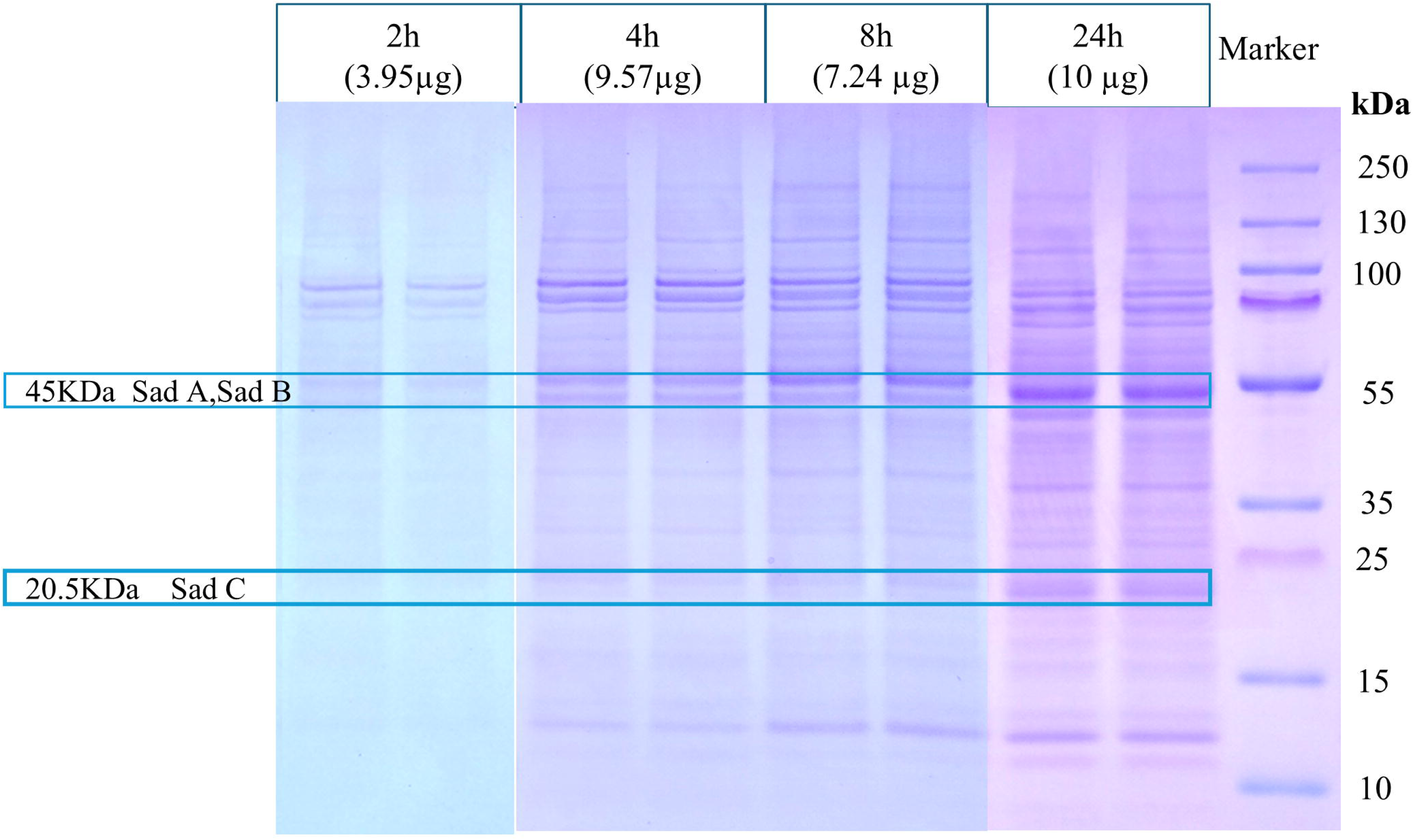
Bis-Tris Nu PAGE gel (4 - 12%) from the test 12 µg L^−1^ SMX stained with Coomasie after an electrophoresis run. The protein extract loaded per lane appears in brackets. Duplicate lanes were loaded per sampling point. The *Marker* on the right column serve as reference for the bands of the samples. Bands aligned to where the Sad cluster enzymes would be retained are enclosed with a frame.

**Figure S2.**
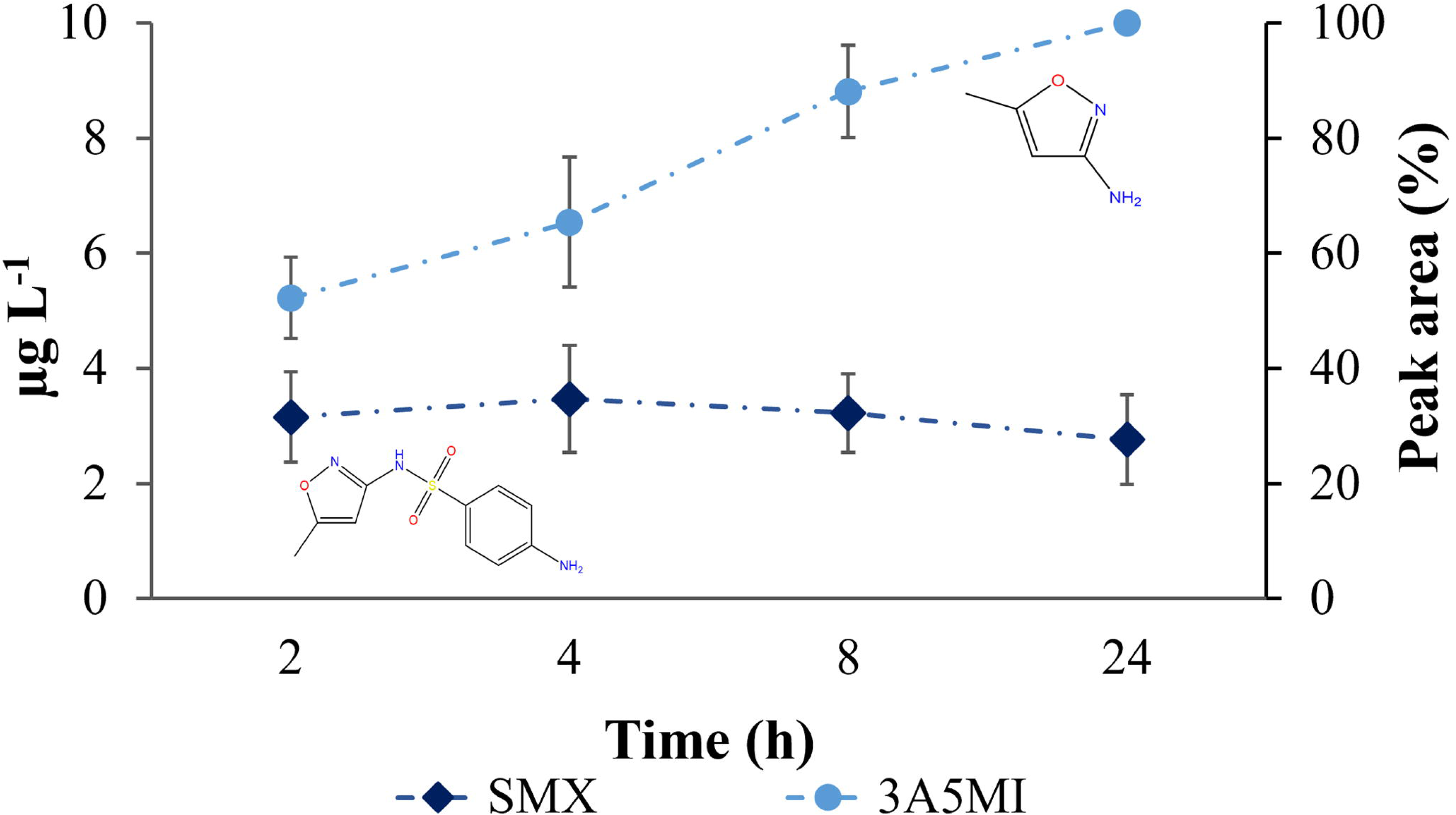
Evolution of SMX (diamond) and 3A5MI (circle) during biotransformation test with an initial concentration of 12 µg L^−1^. 3A5MI corresponds to a relative detected peak area. Depicted values are means of triplicates with their standard deviations.

## Notes

### Competing Interest Statement

The authors have declared no competing interest.

