## Supplemental Material 1 for "Grasping the constraints of pure bacterial strains for the complete catabolism of micropollutants: a proteomic and kinetic study"

Supplementary data contains:

4 Pages

1 Text

2 Tables

2 Figures

**Text S1. Analytical measuremet of the parent SMX and 3A5MI.** Defrost supernatants from the biotransformations tests with concentrations of 20µg L^-1^ and 12 µg L^-1^ were analyzed in the UHPLC ELUTE with a method including OLE up-concentration and QTOF analyzer. A C18 column model Intensity solo (Bruker) was selected for the chromatographic separation of the 100 µL injected volume. The technical especifications of the column include 100 mm lenght, 2.1 µm inner diameter, 2 µm size particle and pore size of 100 Å. For the elution, the mobile phase consisted of A) H_2_O+ Formic acid (FA) 0.1% and B) Methanol + FA 0.1%. with a constant flow rate of 0.250 mL min^-1^and a gradient described in the Table S1. The analyzed mass range comprised from 50 to 750 m/z. A calibration curve prepared in PBS with the analytical standards of SMX and 3A5MI served for the calibration of both compunds. Nevertheless, the ionization of 3A5MI was not optimal at various stages of the run, leading to the analysis of relative units (peak areas). The specific m/z monitored corresponded to 99.055 for 3A5MI with a retention time (RT) of 3.20 minutes and 254.059 for SMX at RT=3.90 minutes. The respective limits of quantification(LOQ) were 0.1 µg L^-1^ for 3A5MI and 0.5 µg L^-1^ for SMX. Mass spectrometry analyses were performed at the Mass Spectrometry and Proteomics Unit (Area of 201 Infrastructures) of the University of Santiago de Compostela.

**Table S1. Flow gradient of the mobile phases used in the UHPLC ELUTE for the analysis of SMX and 3A5MI.**

| Time (min) | %A | %B |
| --- | --- | --- |
| 0 | 95 | 5 |
| 0.4 | 95 | 95 |
| 0.5 | 75 | 25 |
| 4 | 25 | 75 |
| 6 | 0 | 100 |
| 12 | 0 | 100 |
| 12 | 95 | 5 |
| 15 | 95 | 5 |

**%A=** % H_2_O+ FA 0.1%; **%B**= % Methanol + FA 0.1%.

**Table S2. Protein concentration from extracted pellets of *Microbacterium sp* BR1 quantified with the BCA test.**

| Test concentration | Sampling time  (h) | Protein  (µg mL^-1^ )* |
| --- | --- | --- |
| 0.1 µg L^-1^ | 2 | 386.67 |
|  | 4 | 891.67 |
|  | 8 | 690 |
|  | 24 | 1145 |
| 12 µg L^-1^ | 2 | 955 |
|  | 4 | 1106.67 |
|  | 8 | 766.67 |
|  | 24 | 1455 |
| 20 µg L^-1^ | 2 | 388.33 |
|  | 4 | 540 |
|  | 8 | 940 |
|  | 24 | 1386.67 |

All the samples were resuspended in water molecular grade

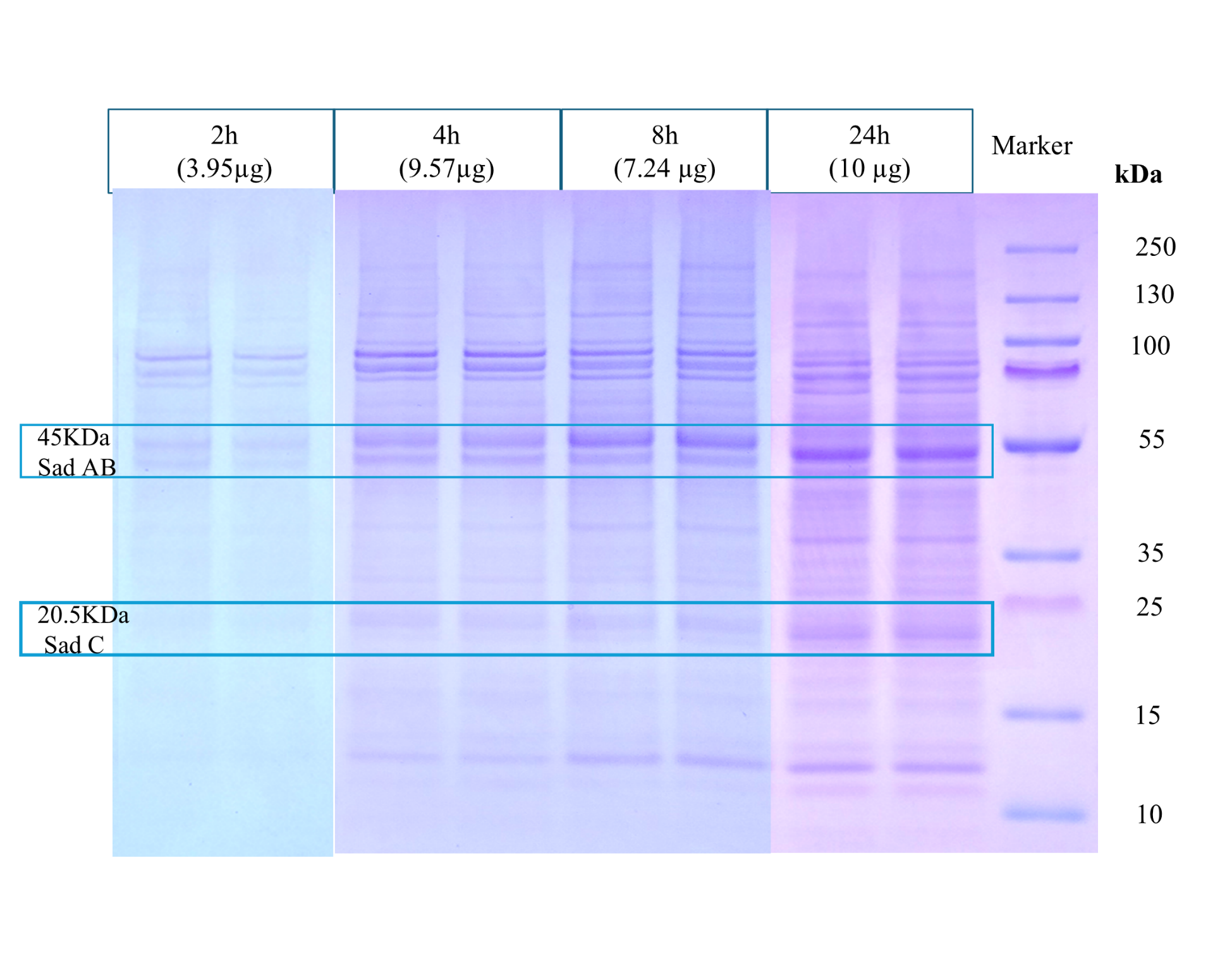
*The injected volume was adjusted per concentration to analyze comparable extracted proteins

**Figure S1. Bis-Tris Nu PAGE gel (4 – 12%) from the test 12 µg L^-1^SMX stained with Coomasie after an electrophoresis run.** The protein extract loaded per lane appears in brackets. Duplicate lanes were loaded per sampling point. The *Marker* on the right column serve as reference for the bands of the samples. Bands aligned to where the Sad cluster enzymes would be retained are enclosed with a frame.

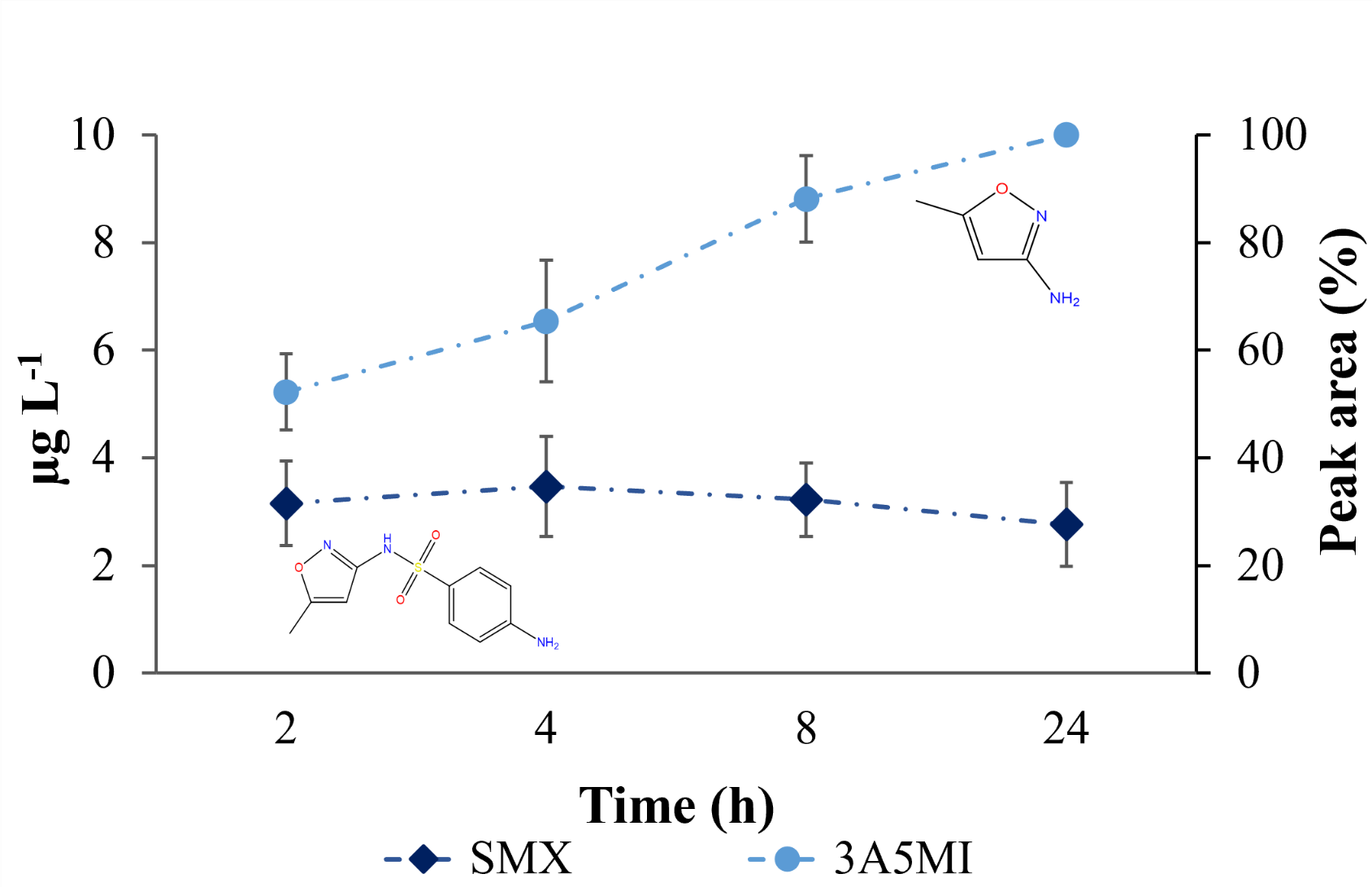
**Figure S2.** Evolution of SMX (diamond) and 3A5MI (circle) during biotransformation test with an initial concentration of 12 µg L^-1^. 3A5MI corresponds to a relative detected peak area. Depicted values are means of triplicates with their standard deviations.
